# Visual Working Memory Guides Attention Rhythmically

**DOI:** 10.1101/2025.06.18.660380

**Authors:** Jiachen Lu, Yaochun Cai, Xilin Zhang

## Abstract

How does internal representation held in visual working memory (VWM), known as the attentional template, guide attention? A longstanding debate concerns whether only one (Single-Item-Template theory) or multiple (Multiple-Item-Template theory) items serve as attentional templates simultaneously. Here we propose a Rhythmic-Item-Template hypothesis, successfully reconciling these seemingly contradictory theories. Using the classical VWM-guided attention task, we found that two VWM items alternately dominate behavioral guidance in theta-rhythmic (4–8 Hz), with anti-correlated activation states in time, and more importantly, this rhythmic oscillation was not driven by the retro-cue processing. Neural recordings revealed that occipital alpha-oscillation (8–14 Hz) governed item-specific prioritization and its amplitude closely tracked subjects’ behavioral guidance, while frontal theta-oscillations phase-led and coupled with occipital alpha-oscillations during the item transition. Our Rhythmic-Item-Template results not only resolve previous Single-Item-Template versus Multiple-Item-Template debate but also advance our understanding of how distributed brain rhythms coordinate flexible resource allocation in multi-item memory systems.

## Introduction

The capacity of visual working memory (VWM) to guide attention through multiple internal templates remains contested^1^. While VWM can store several items^2–4^, behavioral studies conflict: some suggest only one item dominantly guides attention at any moment^5,6^, whereas others report parallel guidance by multiple templates^7,8^. Hollingworth and Beck^9^ revealed that distractors matching either of two target colors elicited equivalent attentional capture, supporting dual templates. However, attenuated capture magnitude compared to single-item conditions exposed a critical paradox: if multiple templates coexist, why does their behavioral efficacy diminish? Three hypotheses emerge: (1) transient dominance of a single item suppresses others, (2) independent but weakened template influences, or (3) rhythmic alternation between items via theta-band oscillations (4 – 8 Hz). Critically, the third hypothesis posits that limited attentional resources are dynamically allocated through cyclical prioritization rather than static competition—a mechanism aligning with the oscillatory nature of neural processes^10,11^.

This oscillatory framework is rooted in attention’s discrete temporal dynamics. When monitoring two locations, human attention rhythmically samples each location at 4–10 Hz, with asynchronous peaks between sites — a “sequential attentional spotlight”^12^. Neural recordings demonstrate that alpha-band suppression (8–14 Hz), reflecting attentional engagement, alternates between objects every ∼200 ms (theta rhythm^13^). These findings imply a conserved theta-rhythmic mechanism for resolving representational competition, whether for external stimuli or internal VWM representations. Recent studies extend this to VWM: multi-item retention exhibits 6 – 10 Hz oscillatory patterns^14–16^, yet debates persist about whether these rhythms reflect true prioritization dynamics or spatiotemporal confounds ^17,18^.

Moreover, Previous evidence for rhythmic memory processing primarily stems from behavioral measures, lacking direct neural evidence of stimulus-specific modulation. While alpha oscillations (8 – 14 Hz) in visual cortex track VWM item prioritization^19–21^, they likely enhance signal-to-noise ratios through selective neural recruitment rather than directly governing task goals^22–24^. Conversely, frontal theta oscillations (4 – 8 Hz) coordinate goal-directed behaviors and task switching^25,26^, exerting top-down control over sensory regions via phase synchronization^27,28^. Critically, fronto-posterior network coupling mediates control shifts in VWM: increased frontal theta power predicts contralateral occipital alpha suppression during priority transitions^29–31^. This supports a mechanistic hypothesis: frontal theta oscillations rhythmically drive spatially distributed alpha oscillations through cross-frequency coupling, dynamically activating multiple VWM items in theta-rhythmic cycles.

To test this theoretical proposition, we designed and conducted three experiments using a memory - search paradigm that effectively rules out strategic control^32^. Experiment 1, with dense temporal sampling, revealed 7 Hz anti - phasic oscillations in attentional capture between two VWM items. Experiment 2, by implementing individualized temporal analysis to eliminate retrospective cueing artifacts, replicated the 7 Hz behavioral rhythm. To further explore the neural mechanisms and comprehensively validate our theoretical hypothesis, we carried out Experiment 3, in which electroencephalography (EEG) was employed to record and analyze brain electrical activity with high temporal resolution. EEG results demonstrated that activation in the occipital alpha band was closely associated with behavioral performance and exhibited significant phase coupling with prefrontal theta oscillations. Additionally, during the memory maintenance phase, the coupling strength of alpha - theta between the bilateral visual cortex and the prefrontal cortex alternated in a theta - rhythmic pattern in leading. These results collectively establish frontally driven theta - alpha coupling as a mechanism for rhythmically allocating attentional access to multiple VWM items, reconciling the divergence between single - and dual - template theories through a dynamic, oscillation - based model.

## Results

### Experiment 1: theta-rhythmic oscillation of visual working memory

25 participants engaged in Experiment 1. As shown in Figure 1C, subjects were asked to remember two memory items that appeared simultaneously. During the memory retention interval, a cue randomly instructed to one of the items, directing internal attention towards the cued stimulus. To examine the time-course of VWM, the response interface appeared at various time intervals following the cue, with a high temporal resolution (stimulus onset asynchrony, SOA: starting at 233 ms and increasing in increments of 33 ms up to 867 ms). The response interface required participants to complete either a search task (80% of trials) or a recall task (20% of trials). In the search task, participants were asked to search a target square (with a gap at the top/bottom) while disregarding a distractor square (with a gap on the left/right). To ensure the search target and the memory item did not share the same spatial location, the target and distractor were positioned vertically relative to the participants’ point of gaze. Only one of the target and distractor matched the color of memory items, or neither did. The attentional capture effect was measured as the difference in response time between the distractor matching the memory items and the target matching the memory items. In the recall task, participants were asked to determine whether a presented color corresponded to one of Initially, we calculated the recall accuracy for both memory items, as depicted in Figure 1B. The recall accuracy for cued items (t(24) = 91.60; p < 0.001, Cohen’s d = 37.39) and uncued items (t(24) = 70.74; p < 0.001, Cohen’s d = 28.88) significantly exceeded the chance level (0.5), demonstrating effective memory retention for both item types. Subsequently, we examined the attentional capture effects associated with these memory items. The analysis revealed that the magnitude of the attentional capture was significantly above zero for both cued items (t(24) = 6.16; p < 0.001, Cohen’s d = 2.51) and uncued items (t(24) = 8.02; p < 0.001, Cohen’s d = 3.27), indicating that both items effectively captured attention. These findings align with previous studies^9^, confirming that human brain is capable of retaining multiple memory items simultaneously, with each item significantly influencing attentional processes.

**Figure 1.**
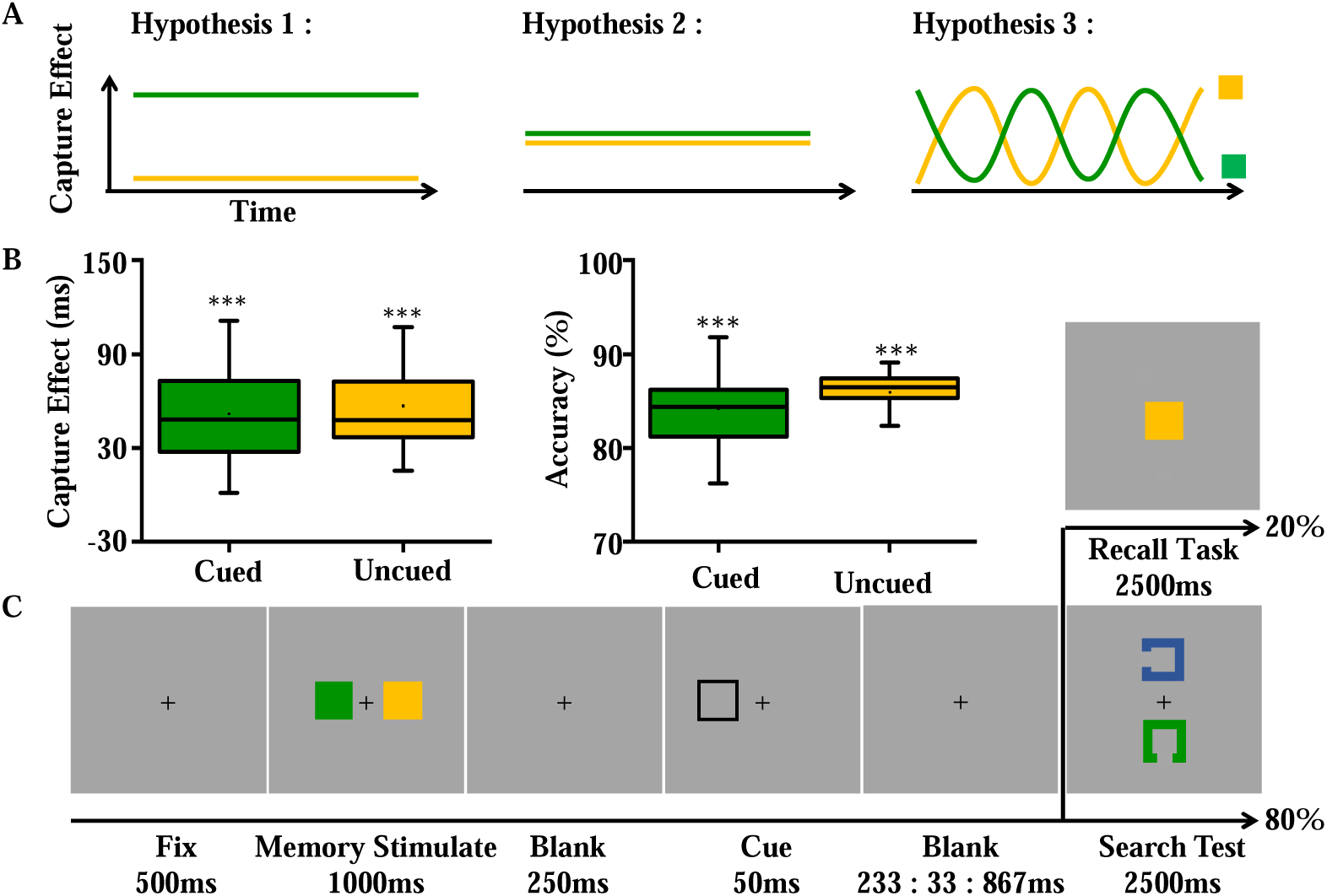
A: Hypothetical models illustrating the activation patterns of two memory items during the memory retention phase. Hypothesis 1 posits a single template, Hypothesis 2 suggests multiple templates, and Hypothesis 3 proposes dynamic templates. B: Behavioral results, showing the attentional capture effect size (calculated as the difference in response time between the distractor matching the memory items and the target matching the memory items) and memory accuracy for the two memory items. C: Experimental procedure. A cue stimulus randomly indicated one of the two memory items. In 80% of the trials, participants performed a search task to identify the item with a gap facing upward or downward. In 20% of the trials, participants performed a recall task to determine whether the probed item matched one of the two memory colors.

With respect to the central question of our study, we found clear evidence for memory item sampling as a preferred template. As shown in Figure 2A, we examined the attentional capture effects of cued and uncued items separately for different temporal SOA conditions. It was found that the capture effects of these two items alternated in dominance rather than remaining stable (F(1, 19) = 4.67, p < 0.001, η2 = 0.16). Specifically, the capture effect of cued items was significantly greater than that of uncued items at SOAs of 267ms (t(24) = 2.72, p = 0.03, Cohen’s d = 1.11), 667ms (t(24) = 2.37, p = 0.03, Cohen’s d = 0.97) and 833ms (t(24) = 3.53, p = 0.002, Cohen’s d = 1.44), while the capture effect of uncued items was significantly greater than that of cued items at SOAs of 333ms (t(24) = 2.97, p = 0.007, Cohen’s d = 1.21), 367ms (t(24) = 2.14, p = 0.04, Cohen’s d = 0.87), 433ms (t(24)= 2.49, p = 0.02, Cohen’s d = 1.02), 467ms (t(24)=2.37, p = 0.03, Cohen’s d = 0.97) and 567ms (t(24)=2.72, p = 0.02, Cohen’s d = 1.11).

**Figure 2.**
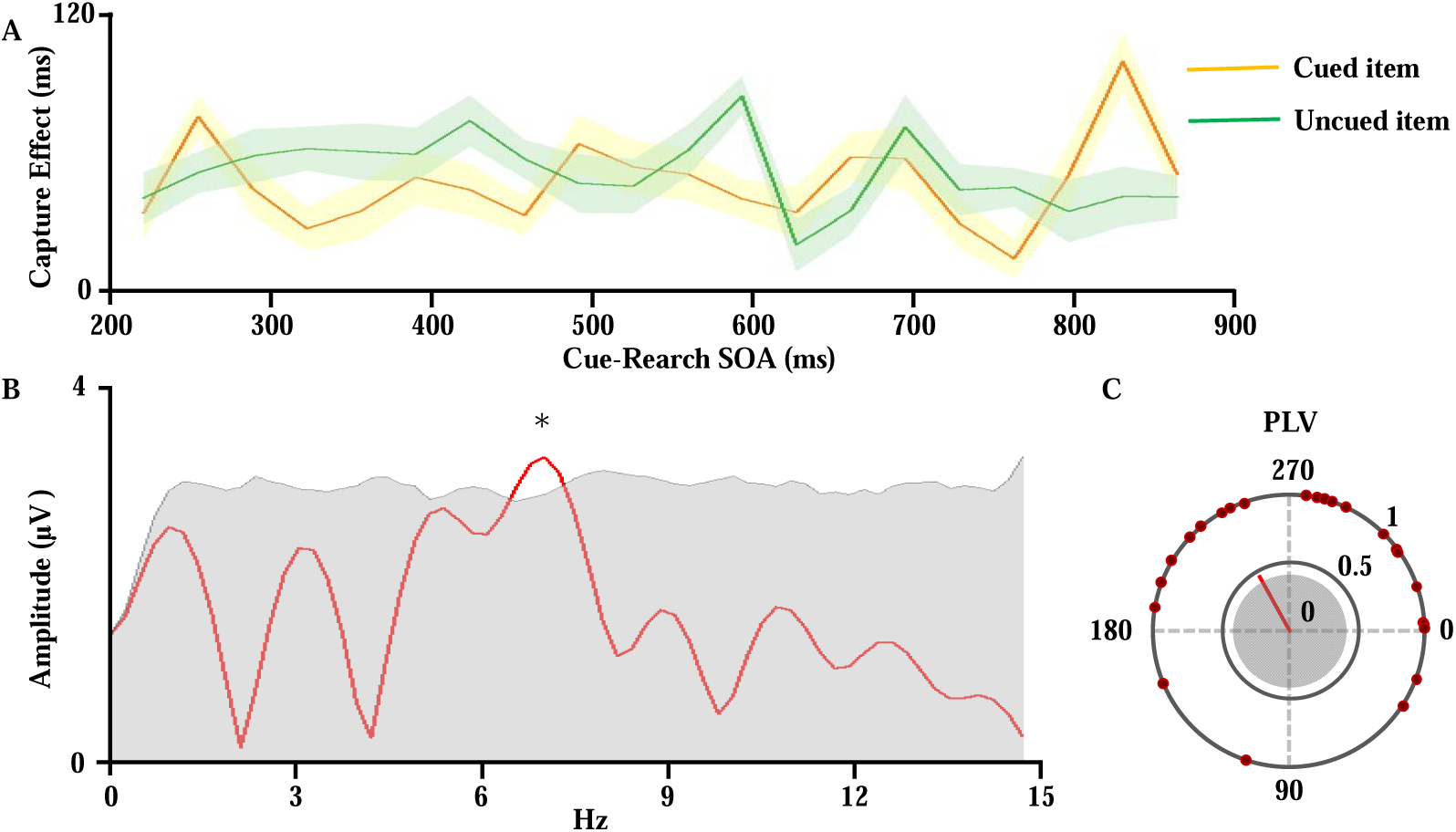
A: Line graph. The attentional capture effect size for the cued and uncued items across different time intervals (SOA), calculated as the difference in response time between the invalid and valid conditions. B: Spectrum plot. The red line represents the amplitude of the values from A at different frequencies; the gray line indicates the 95th percentile from the permutation test; *: p < 0.05. C: Phase-locking value (PLV). The red line shows the PLV values at 7 Hz for the cued and uncued items, representing the average phase difference across all participants; gray circles indicate the 0-95 percentile range from the permutation test; blue hollow circles represent the phase differences for individual participants between the two items.

In addition, as shown in Figure 2B, the Fourier transformation was applied to the time course of the item-based benefit to determine the temporal frequency of these fluctuations. The item-based benefit was calculated as the difference in capture effects between the cued and uncued item for equidistant time intervals ranging from 200 ms to 833 ms. The amplitude spectrum analysis of frequencies ranging from 1 Hz to 15 Hz (Figure 2B) revealed a significant peak at 7 Hz (p < 0.05, FDR corrected). This finding implies that the rhythmic fluctuations in attentional capture associated with VWM are likely driven by an oscillatory mechanism within the theta frequency range, supporting the proposed sampling frequency in VWM.

Consistent with the hypothesis that only a single memory item serves as a prioritized template at any given moment, the speed at which cued and uncued items are accessed should exhibit a negative correlation, as demonstrated in previous research on visuospatial and object-based attention^12,33^. To investigate the possibility that attention within VWM alternates between items, we analyzed the Fourier coefficients at 7 Hz corresponding to the attentional capture effects for both cued and uncued items and subsequently calculated their phase-locking values (PLV^33^). As shown in Figure 2C, the phase-locking between the 7 Hz rhythms of the two items was found to be significant (PLV = 0.402, p < 0 .05), displaying an average phase angle difference of 118° (95% CI from 48° to 188°). This anti-phase relationship between the oscillations supports the notion that internal attention alternately samples items in working memory.

### Experiment 2: Behavioral oscillation without cueing

17 participants took part in Experiment 2, which closely replicated the design of Experiment 1 with one critical modification: no retro-cue was presented after the two memory items. Instead, the probe appeared directly after one of three fixed stimulus-onset asynchronies (SOAs: 233 : 33 : 867 ms; see Figure 3A). This adjustment was implemented to rule out the possibility that working memory oscillations were driven by cue processing rather than intrinsic maintenance mechanisms.

**Figure 3.**
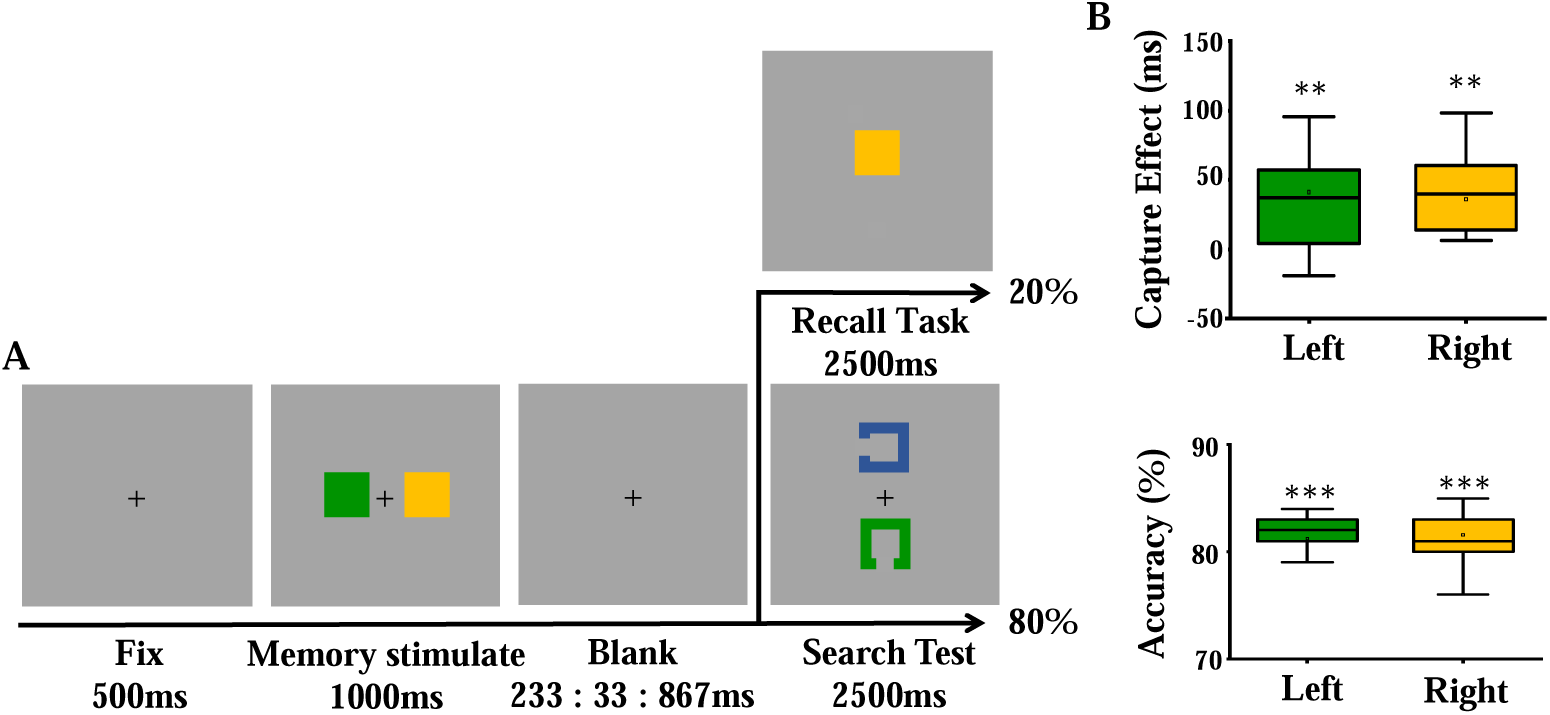
A: Experimental procedure. Two memory items are presented simultaneously without any post-cue prompts. In 80% of the trials, participants performed a search task to identify the item with a gap facing upward or downward. In 20% of the trials, participants performed a recall task to determine whether the probed item matched one of the two memory colors. B: Behavioral results, showing the attentional capture effect size and memory accuracy for the two memory items.

As illustrated in Figure 3B, the recall accuracy for both the left memory item (t(16) = 53.40, p < 0.001, Cohen’s d = 13.35) and the right memory item (t(16) = 51.10, p < 0.001, Cohen’s d = 12.78) significantly exceeded the chance level (0.5), suggesting effective retention of the memory items by participants. Furthermore, our analysis of the attentional capture effect of both memory items revealed that the effect size was notably higher than zero for both the left item (t(16) = 3.24, p = 0.005, Cohen’s d = 0.81) and the right item (t(16) = 3.21, p = 0.005, Cohen’s d = 0.80), demonstrating that both memory items were effective in capturing attention.

Since no retro-cue was presented in Experiment 2, the data lacked a fixed temporal reference point across participants. Thus, unlike in Experiment 1 (where capture effects were group-averaged before Fourier analysis), we performed Fourier transforms individually for each participant before grand-averaging. Specifically, we computed the item-based benefit for each participant by subtracting the capture effect of the right memory item from that of the left item, evaluated at equidistant time intervals (233 – 867 ms). For each participant, we conducted amplitude spectrum analysis (1–15 Hz) on these time series. The resulting spectra were then grand-averaged across participants (blue line, Figure 4). To assess whether spectral power exceeded chance levels, we generated a null distribution by (1) randomly shuffling each participant’s time series, (2) recomputing the Fourier transform, and (3) repeating this procedure 1,000 times. The median amplitude of these surrogate data served as the chance baseline. A paired-samples T-test comparing the original amplitudes against this null distribution revealed significant oscillatory power at 7 Hz (p < 0.05, FDR corrected).

**Figure 4.**
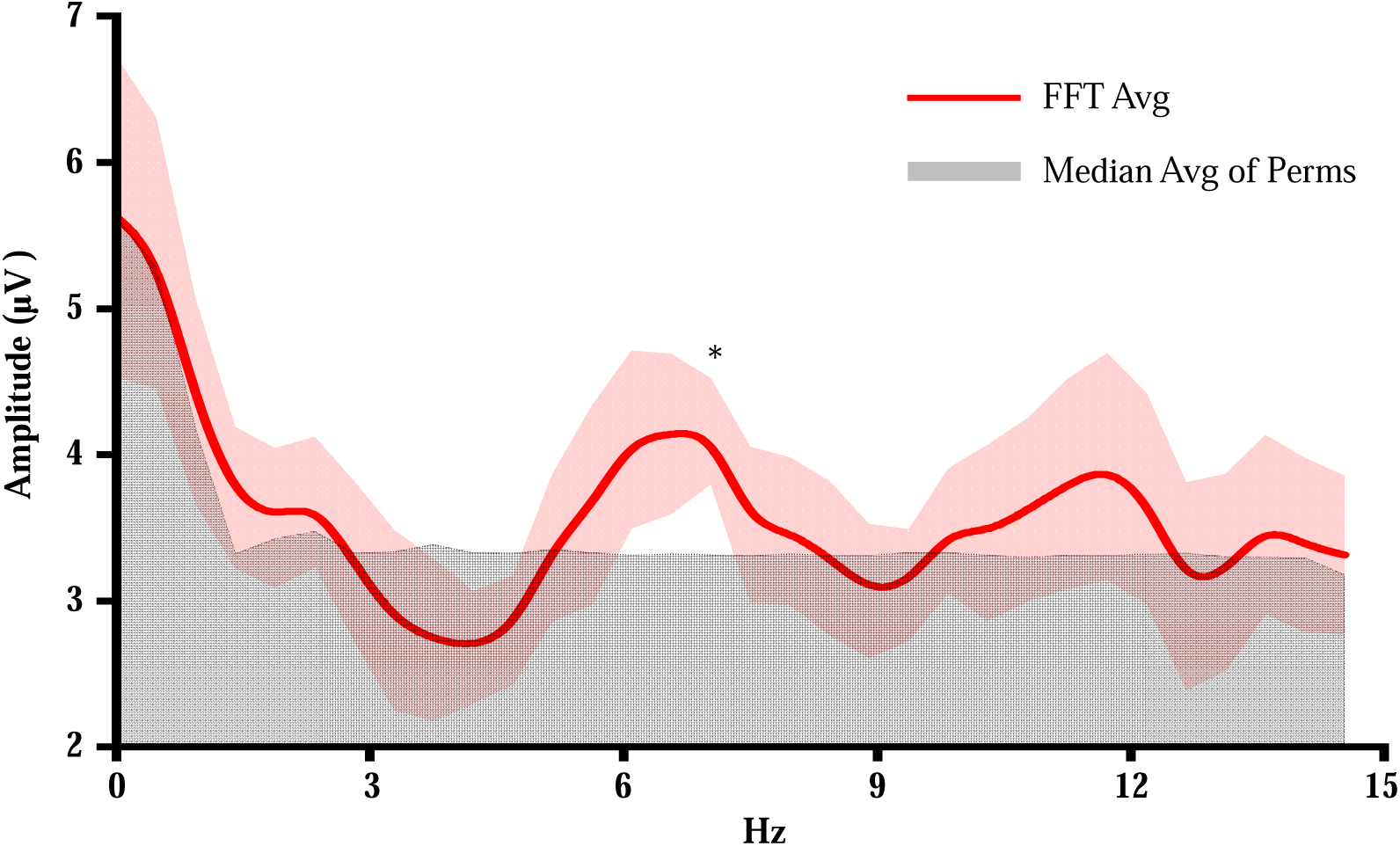
The red line represents the average across all participants of the Fourier transforms of the differences in capture effects between left and right memory items at the individual level. The gray area represents values below the group average of medians derived from 1000 permutations, with each permutation involving Fourier transforms for each participant. *: p < 0.05.

### Experiment 3: Neural mechanisms of visual working memory oscillations

In Experiment 3, 27 participants were recruited, with 3 participants excluded due to excessive artifacts in EEG data. This experiment differed from Experiment 1 in that the search task and the cue were presented at a fixed interval of 1500 ms, while the interval between the memory task and the probe varied randomly between 200 ms and 2000 ms (see Figure 5B). All other aspects of the experiment remained consistent with those of Experiment 1. Additionally, EEG data were recorded simultaneously from participants.

**Figure 5.**
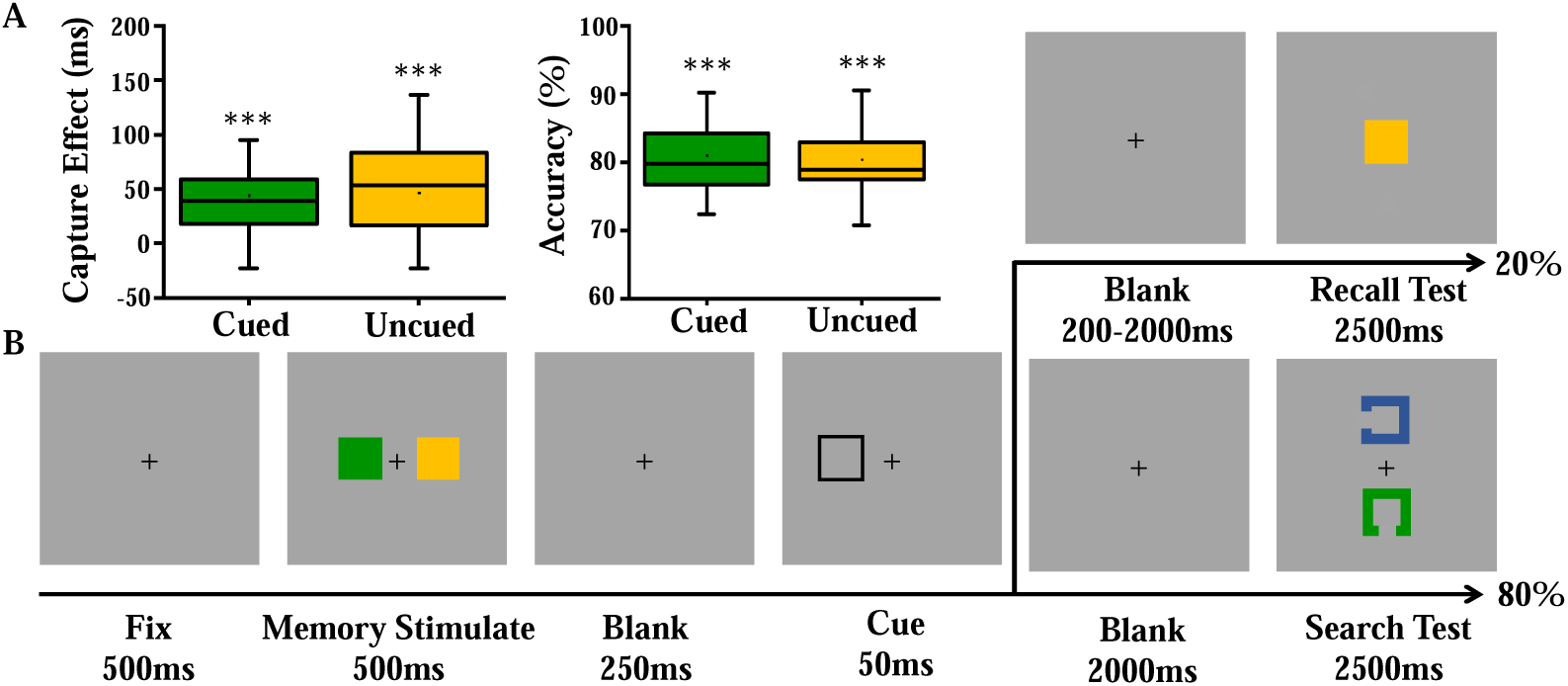
A: Behavioral results, showing the attentional capture effect size and memory accuracy for the two memory items. B: Experimental procedure. A cue stimulus randomly indicated one of the two memory items. In 80% of the trials, participants performed a search task to identify the item with a gap facing upward or downward. In 20% of the trials, participants performed a recall task to determine whether the probed item matched one of the two memory colors.

First, we calculated the recall accuracy for both memory items. As shown in Figure 5A, cued (t(23) = 45.70, p < 0.001, Cohen’s d = 19.06) and uncued items (t(23) = 23.56, p < 0.001, Cohen’s d = 9.82) were both recalled correctly at a significantly greater level than the chance level (0.5), indicating successful retention of both types of items. Additionally, we analyzed the attentional capture effects of the two memory items. The findings revealed that the attentional capture effects for cued (t(23) = 6.45, p < 0.001, Cohen’s d = 2.69) and uncued items (t(23) = 5.61, p < 0.001, Cohen’s d = 2.34) were significantly greater than zero, indicating that both types of memory items effectively captured attention.

Subsequently, an individual time-frequency analysis was conducted on the EEG data during the memory retention phase before the search task. This analysis delved into the temporal and spectral dynamics across frequencies from 1 to 30 Hz and time intervals from −500 to 2000 ms for each participant. Consistent with prior studies^34^ ^19^, pronounced alpha band activity (8 - 14 Hz) was observed in regions contralateral (PO7/8) to the memorized items (see Figure 6).

**Figure 6.**
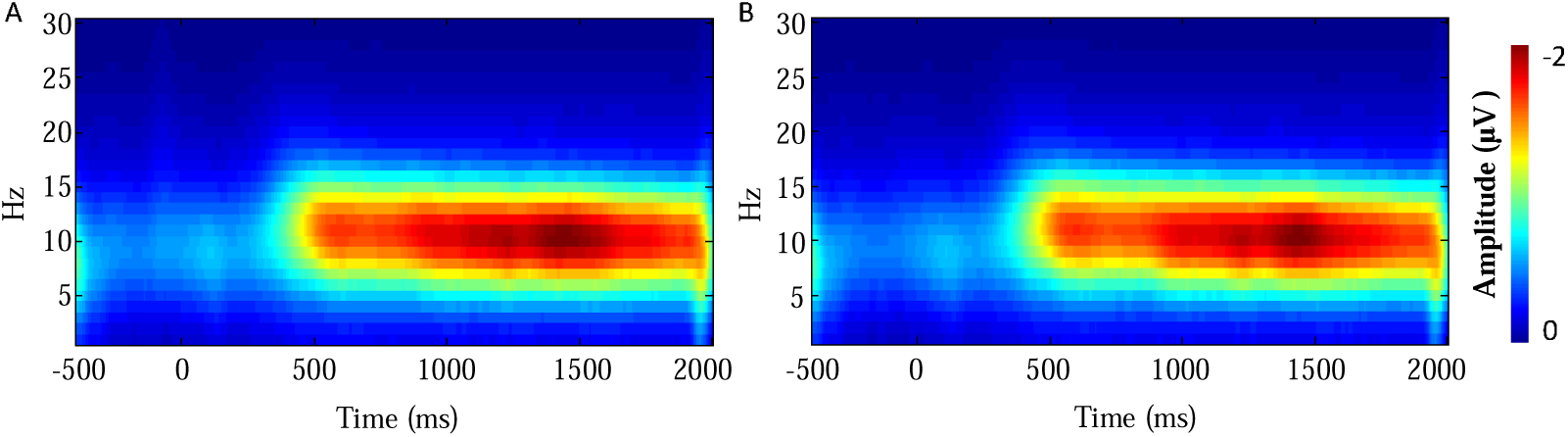
A: Time-frequency results of the contralateral occipital lobe (PO7/8) for cued items, showing significant alpha band activation (8-14 Hz) during the memory retention phase. B: Time-frequency results of the contralateral occipital electrodes (PO7/8) for uncued items, with a similar activation pattern to that observed for cued items.

To evaluate the impact of alpha band activity on attentional capture, correlation coefficients between behavioral performance and alpha power were calculated. Specifically, the correlation between the alpha power contralateral to the items 200 milliseconds before the onset of the search task and the capture effect of the item was analyzed. A full-sample Pearson correlation revealed a significant positive relationship for the cued item (r = 0.43, p = 0.03). To assess robustness, we conducted leave-one-out cross-validation, which yielded positive correlation coefficients across all 24 iterations (range: 0.183 – 0.497, mean r = 0.430 ± 0.055), indicating that no single participant drove the effect. However, the Spearman rank correlation, which is more robust to univariate outliers, was not significant (r = 0.13, p = 0.57), and the p-values from the leave-one-out analysis varied considerably (range: 0.0158 – 0.4025). Taken together, these results suggest a positive trend that is directionally robust but whose statistical significance is sensitive to sample composition. Thus, we interpret this as preliminary evidence that occipital alpha activity may be associated with the priority state within VWM, warranting replication in larger samples. With this caveat, our findings broadly align with prior work linking alpha oscillations to working memory guidance^19,35^.

Analysis of power differences between cued and uncued items unveiled a distinct pattern of alpha inhibition followed by rebound activation within the first 200 ms, as illustrated in Figure 7. Remarkably, this alternation in alpha band power persisted throughout the memory retention phase and showed a significant rhythm of approximately 4 Hz (Figure 9A, p < 0 .05).

**Figure 7.**
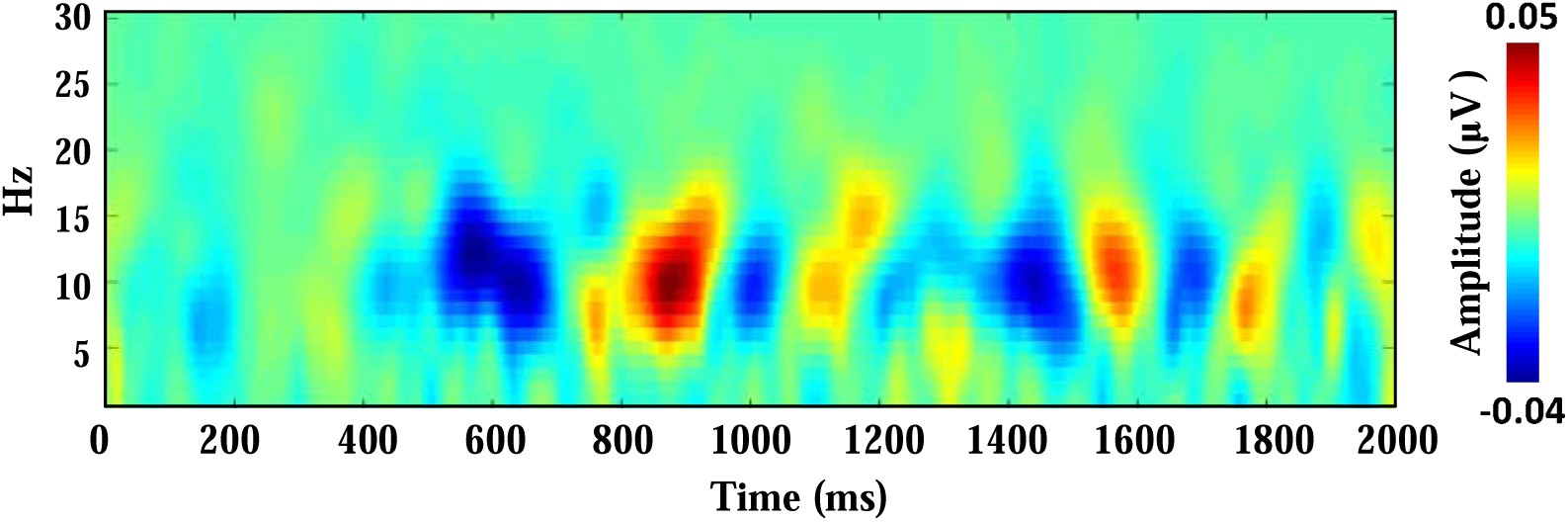
Time-frequency results, showing the difference in contralateral electrode activity (PO7/PO8) for same-color and same-location items under cued and uncued conditions.

Notably, both this study and prior research have identified that the alpha band decodes the preferred memory template. Intriguingly, an observation common to our study and earlier work is that the alternating frequency of this preferred template aligns with the theta rhythm range (4-7 Hz). To explore the potential impact of theta oscillations in different brain regions on alpha oscillations in regions contralateral to the memorized items, we quantified 1:2 cross-frequency phase synchrony (CFS) using the phase-locking value (PLV) between theta (4-7 Hz) and alpha (8-14 Hz) oscillations. Specifically, for each electrode pair we computed 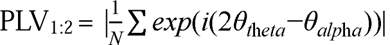 following established methodologies (Siebenhühner et al., 2020; Palva et al., 2005). The findings, illustrated in Figure 8A, indicated significant coupling between PO7/8 and the prefrontal regions. Additionally, CFS analysis revealed significant coupling for theta oscillations in the prefrontal cortex (Fz) and alpha oscillations in the visual regions. Fascinatingly, illustrated in Figure 8B, by assessing the alpha-theta coupling strength between the posterior regions contralateral to the two items (PO7/8) and the prefrontal regions (Fz) on a moment-to-moment basis during the memory retention period, we found that the coupling strength of the two items alternated in lead. To further define the temporal frequency characteristics of these alternations, Fourier transformation was applied to the differences in PLV associated with the two items. An amplitude spectrum analysis across frequencies from 1 Hz to 50 Hz (Figure 9B) identified a significant peak at 4 Hz (p < 0.05, FDR corrected), underscoring the rhythmic interconnection between these neural activities (see Figure 9B).

**Figure 8.**
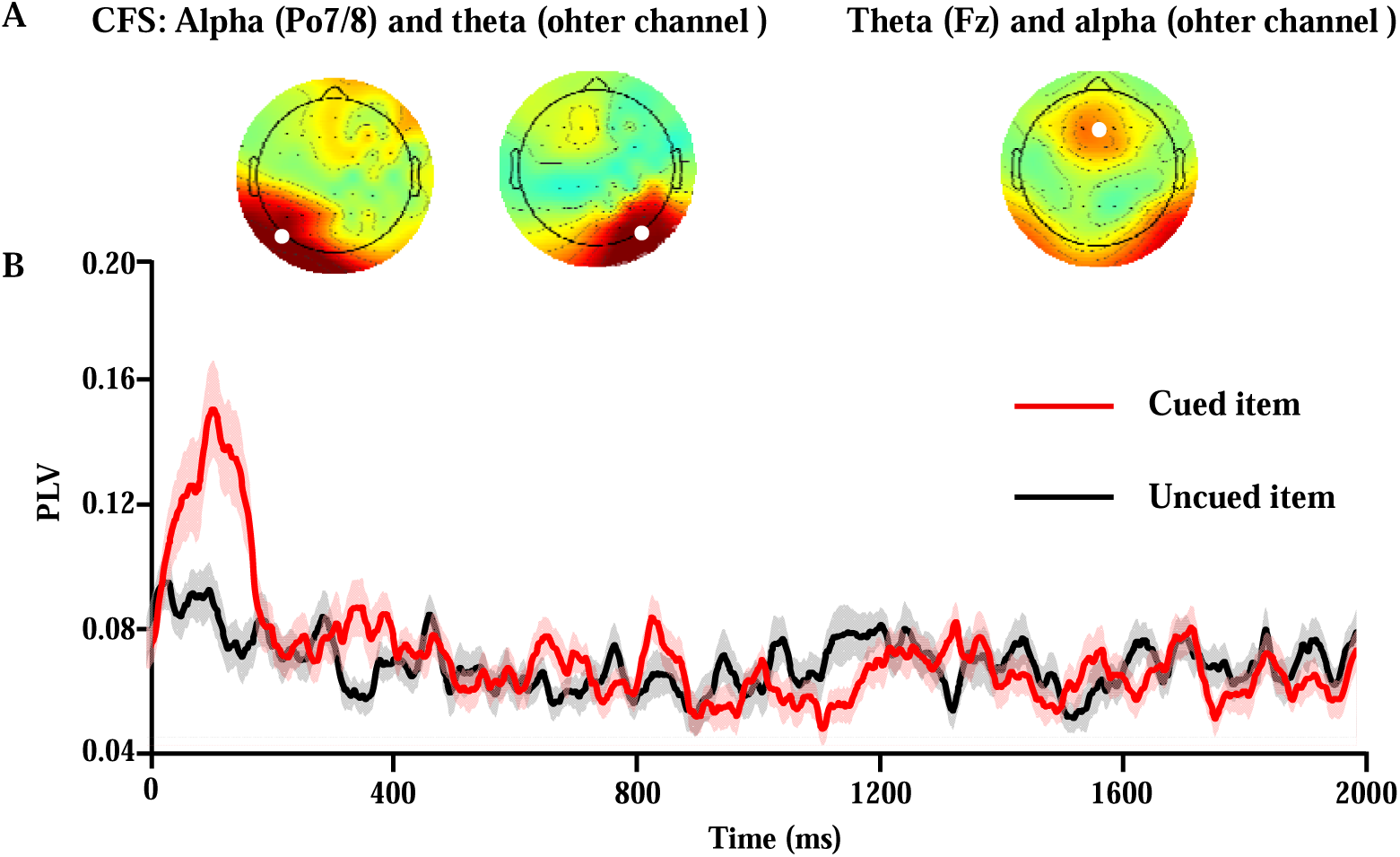
A: Topographical maps. 1:2 cross – frequency phase synchrony (CFS) between the alpha phase at the white electrode points and the theta phase at other electrode points (the left two maps), and the theta phase at the white electrode points and the alpha phase at other electrode points (the right map) during the memory retention phase. B: Functional connectivity map. 1:2 CFS phase coupling between the alpha phase at the contralateral electrodes (PO7/PO8) of the two memory items (red lines represent the cued item; black lines represent the uncued item) and the theta phase at the frontal electrode (Pz) during the memory retention phase.

**Figure 9.**
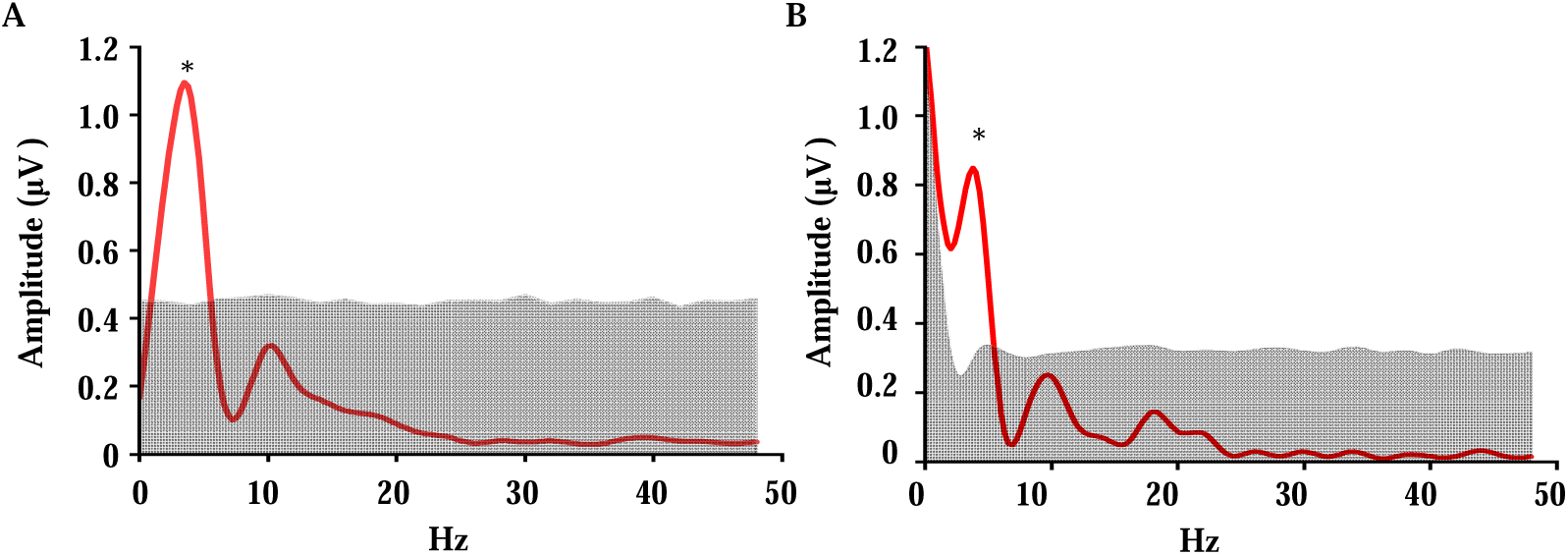
A: Spectrum results. Power spectrum of the difference in contralateral alpha power between the two memory items during the retention phase across 0-50 Hz. B: Spectrum results. Power spectrum of the difference in cross-frequency coupling (PLV-based 1:2 CFS) between the alpha phase at contralateral electrodes (PO7/PO8) and the theta phase at the frontal electrode (Fz) across 0-50 Hz during the retention phase. The gray shaded area represents the 0-95th percentile range from the permutation test; *: p < 0.05.

## Discussion

This study employed the function of VWM as a preferred template to automatically capture attention and integrated EEG techniques to systematically investigate the mechanisms underlying the representation of two memory items that occurred simultaneously or in close temporal proximity during the memory retention phase. Our findings revealed several key insights. Firstly, we observed that the ability of the memory items to capture attention alternated over time, displaying opposite trends (Experiment 1). This indicates that the preference for the memory items alternated rather than occurring simultaneously. Power analysis showed alternating rhythms within the 4-7 Hz range (Experiment 1-3), corresponding to theta oscillations. Secondly, during the memory retention phase, the strength of alpha band oscillations, induced by cued and uncued items, alternated in dominance throughout the retention period. Thirdly, we observed strong rhythmic alternations in the coupling strength between the alpha band of both posterior regions and the theta band of the prefrontal region. The coupling exhibited a leading frequency of approximately 4 Hz. In conclusion, our findings suggest that memory is maintained by alternately processing multiple items within the theta rhythm, rather than continuously entangling a single object. Furthermore, the posterior alpha oscillations reflect the prioritization of memory items and exhibit a close association with theta oscillations in the prefrontal cortex.

Traditional research on working memory has often adopted a static viewpoint, positing that its performance remains constant throughout the retention phase. This perspective, however, is challenged by the findings of the present study, which investigates the ability of two memory items to capture attention under various Stimulus-Onset Asynchrony (SOA) conditions. Interestingly, the study reveals that the attentional capture abilities of these memory items alternate in dominance. Spectral analysis of the SOA function identified significant rhythmic fluctuations at 7 Hz, with opposing trends. This pattern suggests that, in scenarios requiring the maintenance of two working memory items for a task, these items alternately serve as the dominant template during the retention phase.

This discovery aligns with the existing literature on the rhythmic processing of external stimuli. Specifically, it has been well-documented that attention oscillates within the theta rhythm (4-8 Hz) across various task types, including those based on spatial attention ^36,37^, feature attention^38^, and object attention^10,33,39^. Moreover, rhythmic attentional modulation has been observed in auditory attention^40,41^, in the effects of top-down predictions on brightness perception^42^, and in the process of visual feature binding^43^. These findings collectively highlight that attention operates within a theta rhythm, extending beyond external stimuli to include the internal processing of memory information^15,16^.

Similar to the present study, Peters et al. ^15^ had participants memorize four spatial positions forming the endpoints of two objects (one cued), and their results showed that after the two objects disappeared, attention fluctuated at the theta rhythm between their original positions with an inverse correlation; in contrast, the present study explores the manner of memory maintenance indirectly by leveraging the guiding effect of working memory on attention, effectively avoiding the influence of spatial positions—while Peters et al.’s study, which directly examined differences in probe positions, clearly demonstrates that attention undergoes rhythmic changes at the two spatial locations and persists after the objects vanish, it hardly clarifies the rhythmicity of working memory performance, whereas the present study directly investigates such performance using the attention-capture effect of working memory, revealing that when maintaining multiple memory items, their attention-capturing capabilities alternate in dominance, i.e., multiple working memory items alternately become priority templates in a rhythmic manner.

The concept of a dynamic template mechanism emerges as a novel resolution to the ongoing debate between the single-template and multi-template hypotheses in working memory research^1,6,44^. his mechanism proposes that when maintaining multiple memory items, an individual represents these items through a single, dynamically shifting template, with each moment guided by a single item—a process that is spontaneous and not triggered by a retro-cue prompting a specific memory item. As observed in my Study 2, even in the presence of retro-cues, multiple memory items still begin with a particular item (consistent with individual habits) and sequentially become priority templates at the theta rhythm. This flexible approach not only aligns with the observed phenomenon of multiple memory items capturing attention, but also provides a new perspective on the discourse on how working memory operates, bridging a critical gap in the existing literature.

The current study reveals that during the memory retention phase, the dominance of posterior lateralized alpha oscillation alternates between two memory items and significantly correlates with behavioral outcomes. This finding aligns with extensive evidence suggesting that alpha oscillations are crucial in modulating visual processing, going beyond mere sensory involvement to encompass functional roles in sensory gating and top-down control of preparatory activity^23,36,45,46^. Notably, alpha band activity modulation is essential for prioritizing both incoming sensory data and stored memory content^47,48^. A seminal example is a study in which subjects memorized two sequentially presented items, demonstrating the capability of posterior lateralized alpha oscillations to differentiate the memory item relevant to the current task, thereby elucidating the role of oscillations in managing the priority of concurrent task-relevant items ^35^.

It’s critical to acknowledge that the investigation of posterior alpha oscillations in working memory, including the design of Experiment 3 in this study’s, often involves spatial differentiation of items^35^. Such spatial segregation allows alpha power modulations to be attributed to specific items through lateralization, achieved by presenting items in distinct hemifields. This methodological choice, driven by the spatial specificity of neurons encoding working memory items, serves as a pragmatic experimental strategy rather than implying that memory item prioritization solely depends on alpha oscillation lateralization^49,50^. For example, de Vries et al.^19^ discovered that posterior, rather than lateralized, alpha oscillations could delineate memory items related to forthcoming tasks.

The findings of this study, revealing strong phase coupling between posterior alpha oscillations and theta oscillations in the prefrontal lobe, contribute to the expanding literature on the intricate interplay between different brain oscillation frequencies and their roles in cognitive functions. The association of prioritization processes with theta oscillations in the frontal cortex is well-documented^30,35,51,52^, underscoring the critical role of frontal cortex in orchestrating goal-directed behavior^53^, managing multiple objectives^54^, and facilitating task switching^55^.

Particularly during working memory tasks, the frontal cortices play a critical role in processing and maintaining abstract goal-related representations and task-specific information. This is supported by the evidence of mixed selectivity in frontal neurons^56^ and their coordination of sensory areas based on this information^57,58^. For instance, fMRI studies have highlighted enhanced connectivity between frontal regions and posterior task-related sensory areas specific to working memory representations and those prioritized by maintenance cues (retrospective cues^59,60^), with this increased connectivity correlating with improved performance of cued memory representations^46,61^.

Such evidence underscores the ability of the frontal cortex to selectively coordinate the activation of visual cortical working memory representations relevant to the current task. As outlined in a review by de Vries et al.^30^, alpha modulation facilitates the flexible tracking of changes in the state of VWM. This top-down control over the current state of VWM is accomplished through prefrontal theta oscillations, which in turn orchestrate alpha oscillations via a broad spectrum of cross-frequency interactions, highlighting a sophisticated mechanism of cognitive control and memory prioritization.

Although the present study successfully revealed the phenomenon of dynamic attentional templates through high-density behavioral sampling and preliminarily explored the underlying mechanisms using high-temporal-precision EEG technology—specifically, that prefrontal theta oscillations and visual cortical alpha oscillations play indispensable roles in this process—there are still some limitations. First, despite its high temporal resolution, the EEG technique used in this study makes it difficult to precisely localize the brain regions influencing dynamic templates due to the brain’s volume conduction effect^62^. Second, the sample size of the current study is small, which may render the results vulnerable to interference from extreme cases^63^. Additionally, to assess cross-frequency interactions we used 1:2 cross-frequency phase synchrony (CFS) quantified by the phase-locking value (PLV)^64–66^. This measure directly captures the stability of the harmonic phase relationship (2θ_theta_ - θ_alpha_) and is robust against volume conduction because the two frequencies are distinct. The PLV values of the two items alternately dominated over time, consistent with the pattern of behavioral data, which is hardly explicable by coincidence. Future research could employ more precise techniques (e.g., MEG devices) to accurately capture activities in different brain regions while maintaining high temporal precision. Furthermore, rhythmic transcranial magnetic stimulation (TMS) could be used to modulate rhythms in various brain regions, altering the oscillation frequencies of working memory to provide causal evidence for this phenomenon.

Another limitation concerns the brain-behavior correlation between alpha power and attentional capture. Although the full-sample Pearson correlation was significant and leave-one-out analyses confirmed a consistently positive direction, the Spearman correlation was not significant and the p-values of leave-one-out iterations varied considerably. This pattern likely reflects the modest sample size (N=24) and a moderate true effect size. Accordingly, the observed correlation should be considered preliminary, and future studies with larger samples are required to confirm whether alpha power reliably predicts the magnitude of working-memory-guided attentional capture.

In conclusion, our study aligns with the evolving understanding that oscillatory neural mechanisms underpin the complex interplay between attention and working memory. It builds on the premise that external attentional processes are intricately linked with neural oscillations, while also highlighting the crucial role of internal attention in modulating working memory representations through rhythmic activity. Our findings support a framework in which internal attention is mediated by a dynamic interaction between prefrontal theta oscillations, serving as a top-down control mechanism, and posterior alpha oscillations, which are instrumental in the selective inhibition or facilitation of memory items. This synthesis not only challenges conventional static views of working memory but also proposes a refined model of cognitive processing, where memory maintenance and attentional prioritization are orchestrated within a rhythmic neural symphony.

## Materials and methods

### Participants

A total of 25 participants (9 males, aged 22 ± 1.9 years) took part in Experiment 1, 17 subjects (6 males, aged 19 ± 1.5 years) participated in Experiment 2, and 27 subjects (12 males, aged 21 ± 2.0 years) were involved in Experiment 3. All participants had normal or corrected-to-normal vision and no history of psychiatric or neurological disorders. The experiments were conducted in accordance with the Declaration of Helsinki and received ethical approval from the Research Ethics Committee at South China Normal University (approval date: 2021-04-01). Before the start of the experiment, all participants provided written informed consent. Participants completed the experiment in a dark, quiet and isolated room, with their heads fixed on a head rest and their eyes looking directly at the centre of the screen at a distance of 57 cm from the display.

### Stimuli

The same stimuli were used for all three experiments. The memory stimuli were a square in 6 colours: cyan (RGB: 5, 200, 200), red (RGB: 200, 80, 40), yellow (RGB: 200, 200, 5), blue (RGB: 5, 200, 200), purple (RGB: 112, 48, 160), and green (RGB: 50, 200, 50), each with a visual angle of 1.2 × 1.2°. The search stimuli were colored outlined squares (1.2 × 1.2°, 0.2° line thickness), with a 1.2° gap on the top, bottom, left side, or right side. The color of the search stimulus was consistent with that of the memory stimulus. The cue stimulus is a black square with side length 1.2°. The shape and color of the detection stimulus are consistent with those of the memory stimuli.

The experiments were programmed using the Psychtoolbox in Matlab 2019b. All stimuli were presented on a 17-inch CRT monitor with a resolution of 1024 × 768, a refresh rate of 60 Hz, and a gray background (RGB: 128, 128, 128).

### Tasks

#### Experiment 1

Each trial began with a 500 ms fixation point, followed by a 1000 ms memory array. In the memory array, two differently colored memory stimuli were presented to the left and right at a distance of 3° from the centre of the screen. After a 250ms fixation point, the cue array was presented for 50 ms, with the cue appearing randomly on the left or right at a distance of 3° from the centre of the screen. After a pseudo-random SOA (233 ms : 33 ms : 867 ms), either the search array (80% of trials) or the memory detection array (20% of trials) was presented for 2500 ms. In the search array, two search stimuli were presented directly above or below the screen center, 6° away from it; one had a gap facing up or down (the target), and the other had a gap facing left or right (the distractor). Participants were instructed to respond by pressing the “A” key (for an upward-facing gap) or the “Z” key (for a downward-facing gap). In the memory detection array, a detection stimulus was presented at the screen center for 2500 ms. Participants were asked to press the “N” key (if it belonged) or the “M” key (if it did not belong) to determine whether the detection stimulus belonged to either of the memory stimuli in the memory array, regardless of the cue. After responding, participants moved on to the next trial, with a 1000ms interval between trials. To familiarize participants with the task, 15 practice trials were conducted before the formal experiment. Participants completed 30 blocks over four days within one week, with each block containing 150 trials (120 trials for search tasks and 30 trials for memory detection tasks).

#### Experiment 2

Each trial began with a 500 ms fixation point, followed by a 1000 ms memory array. In the memory array, two memory stimuli of different colors were presented on the left and right sides, 3° away from the center of the screen. After a pseudo-random SOA (233 ms : 33 ms : 867 ms), either the search array (80% of trials) or the memory detection array (20% of trials) was presented for 2500 ms. Participants’ tasks in the search array and memory detection array were consistent with those in Experiment 1. Over the course of one week (completed across four days), participants finished 30 blocks, with each block consisting of 150 trials (120 for search tasks and 30 for memory detection tasks).

#### Experiment 3

In Experiment 3, high temporal precision EEG technology was employed, which eliminated the need for dense SOA manipulation when investigating dynamic features. Additionally, to enable a longer memory retention time window during analysis, the interval between the cue array and the search array was fixed at 2000 ms. Furthermore, to encourage active maintenance of memory content during the retention phase, the interval between the cue array and the memory detection array was randomly varied between 200 ms and 2000 ms. Consistent with Experiment 1, 20% of the trials involved the memory detection array, and 80% involved the search array. The tasks for both the memory detection array and the search array were the same as in Experiment 1, and both were presented for 2500 ms. Participants were required to complete 4 blocks, with each block consisting of 150 trials.

### EEG recordings and preprocessing

EEG data was recorded using a cap with 64 electrodes arranged according to the international 10–20 system (Brain Products, Munich, Germany). The frontal electrode FCz was utilized as the online reference point, and the AFz electrode was employed as the ground. All electrodes were amplified using a 0.01–70 Hz online band-pass filter and continuously sampled at a rate of 1000 Hz per channel.

The offline continuous EEG data was preprocessed using EEGLAB, an open-source toolbox within the MATLAB environment. Initially, re-reference was conducted by using the bilateral mastoids TP9 and TP10. Next, all EEG signals underwent a 0.1 Hz high-pass filter, a 30 Hz low-pass filter, and a 50 Hz notch filter. Subsequently, independent component analysis (ICA) was applied to each participant;s data to eliminate components related to eye movements and artifacts. The remaining components after this process were then projected back into the channel space. We extracted data from −500 ms to 2000 ms relative to cue stimulus presentation in Experiment 3.

### Data analysis

#### Behavioral performance analysis

We utilized the difference in reaction times (ΔRT) between the invalid trials (where the color of the memory item matched that of the interfering item) and the valid trials (where the color of the memory item matched that of the target item) extracted from the search array as the capture effects.

Subsequently, we employed a one-sample t-test to separately examine whether the memory accuracy rates of the two memory items in the three experiments were significantly higher than the guessing level (50%), and whether the capture effects of the two memory items were significantly greater than 0. In this process, the memory accuracy rates and capture effects in Experiment 1 and Experiment 2 were the results of averaging all the SOA (stimulus onset asynchrony) conditions.

For Experiment 1, we concentrated on exploring how the capture effect of the two memory items on attention evolved over time. To accomplish this, we carried out a repeated-measures analysis of variance (ANOVA) with a 2 (cued item vs. uncued item) × 20 (all SOAs) design. Post-hoc comparisons were conducted using paired-sample t-tests to thoroughly investigate the potential changes in the capture effect of the memory items on attention.

In order to obtain the frequency spectrum of memory-based attention capture over time, we analyzed the difference in capture effects between the cued and uncued items across all SOAs at evenly spaced temporal intervals. Subsequently, we performed a fast Fourier transform (FFT) to estimate the spectral composition, which yielded power values across 14 frequency bins ranging from 1 Hz to 14 Hz. Regarding the phase relationship of 7-Hz oscillations between the cued and uncued items, we conducted separate Fourier transforms for these conditions as previously described. Then, we calculated the angular difference between the phase angles of the 7-Hz oscillations for each condition. This angular difference was projected onto the unit circle in the complex plane and averaged across all participants. The length and angle of the resulting vector represented the phase-locking value (PLV^67^) and the mean phase difference. To evaluate the statistical significance, a non-parametric approach was adopted to estimate the probability of the observed data under the null hypothesis. In 1000 permutation samples, each time-course was shuffled before the analysis, generating one mean amplitude spectrum for the memory-based capture effect, and one mean phase difference between the cued and uncued conditions for each permutation sample. To control the false discovery rate at 5%, the individual frequency p-values in the amplitude spectrum were adjusted according to the number of frequency bins, following the method of Benjamini and Hochberg^68^.

For Experiment 2, since Experiment 2 adopted a modified approach due to the absence of retro-cues. In this experiment, we compared the left and right memory items instead of the cued/uncued items. The item-based attentional benefit was calculated for each participant by subtracting the right-item capture effect from the left-item effect at equidistant intervals (233-867 ms). Individual FFTs (1-15 Hz) were performed on these difference scores before conducting a grand average across all participants. To determine the significance thresholds, we generated null distributions by randomly shuffling each participant’s time series 1000 times, recomputing the FFTs for each permutation, and using the median amplitude as the chance baseline. A paired-samples t-test comparing the original amplitudes against this the chance baseline revealed significant 7-Hz oscillatory power (p < 0.05, FDR corrected), as depicted in Figure 4.

#### Time-frequency decomposition

We used the short-time Fourier transform (STFT) function in Matlab 2019b to Fourier transform the baseline-corrected segments to obtain the power information, where the frequency was set to 30 frequency points of equal length from 1 to 30 Hz and the sliding window was a 200 ms hanning window. The frequency-specific power at each time point was calculated as the square of the amplitude of the complex signal resulting from the convolution, determined by the sum of the squares of the real and imaginary parts^69^. Based on findings from previous studies indicating that memory stimuli can be characterized by alpha oscillations in the contralateral occipital lobes^19^, we therefore extracted alpha band data (8-14 Hz) from PO7 and PO8. We separately averaged the power of the contralateral side for the cued and uncued items. As illustrated in Figure 8, to compare the power induced by cued items with that induced by uncued items during the memory retention phase, we extracted the contralateral power for each color during memory retention. The power when cued was subtracted from the power when uncued. Finally, we averaged the results across all memory colors and participants.

#### Interregional connectivity

We quantified 1:2 cross-frequency phase synchrony (CFS) using the phase-locking value (PLV) between theta (4–7 Hz) and alpha (8–14 Hz) oscillations, following established procedures^64, 65^. Specifically, for each pair of electrodes, we computed:

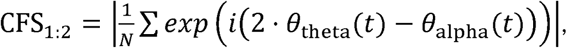

where theta θ_theta_and θ_alpha_ alpha are the instantaneous phases of theta and alpha oscillations, respectively. The factor 2 accounts for the 1:2 frequency ratio (theta : alpha = 1 : 2), effectively aligning the faster alpha oscillation to the slower theta rhythm (phase acceleration). The analysis first revealed that both electrodes PO7 and PO8 exhibited the strongest 1:2 CFS with electrodes in the prefrontal region. Next, we calculated the 1:2 CFS between theta oscillations at the frontal electrode Fz and alpha oscillations at all other electrodes to confirm the specificity of cross-frequency coupling between prefrontal cortex and visual cortex. Finally, we extracted the 1:2 CFS values between the contralateral visual electrodes PO7/8 (relative to the cued and uncued items, respectively) and the frontal electrode Fz.

#### The correlation between neural activation and behavior

As shown in Figure 6, a clear activation of the alpha band was observed during the memory retention phase. To investigate whether this activation was related to behavioral performance, we calculated the alpha activation strength in the 200 ms preceding the search task. Specifically, we computed the average amplitude within the 8-14 Hz frequency range during the 1800-2000 ms time window of the retention phase, contralateral to the cue. This was then correlated with the capture effect size of the cued items using Pearson ’ s correlation. To assess the robustness of this relationship against potential outliers and sample composition, we additionally performed leave-one-out cross-validation and Spearman rank correlation. These complementary analyses are reported in the Results section.

### Frequency spectrum analysis

As shown in Figure 8, the alpha power (8-14 Hz) induced by cued and uncued items alternated in dominance during the memory retention phase. To quantify this rhythmic alternation, we conducted a spectral analysis following these steps: First, we computed the power difference between cued and uncued items within the 8-14 Hz range during the retention phase. These differences were then downsampled to 100 Hz using a 10 ms window for averaging, generating a one-dimensional time series spanning the 0-2000 ms retention period. This time series was subsequently subjected to amplitude spectrum analysis across frequencies from 1 Hz to 50 Hz using Fourier transformation.

To assess the statistical significance of the observed spectral features, we employed a permutation test. Specifically, we randomly shuffled the temporal order of the time series of power differences between cued and uncued items—thereby preserving the amplitude distribution of the data while eliminating temporal correlations in the original sequence—and repeated the Fourier transform and spectral analysis for each shuffled time series. This permutation process was replicated 1000 times to generate a null distribution of spectral power values^70, 71^. A frequency component in the original data was considered statistically significant if its power ranked within the top 5% of the corresponding null distribution (p < 0.05).

We applied the same analytical pipeline to investigate differences in the phase-locking value (PLV) between the contralateral regions of the two items and the prefrontal cortex during the retention phase. Specifically, PLV differences (i.e., the difference between the two conditions) were computed, downsampled to 100 Hz using a 10 ms window for averaging to generate a time series spanning 0-2000 ms, and then subjected to amplitude spectrum analysis (1-50 Hz) using Fourier transformation. Significance was assessed via the identical permutation test procedure described above (randomly shuffling the temporal order of the difference time series).

## Acknowledgements

We acknowledge the subjects for their contribution to this study. This work was supported by National Natural Science Foundation of China (32271099), Research Center for Brain Cognition and Human Development of Guangdong Province (2024B0303390003), Striving for the First-Class, Improving Weak Links and Highlighting Features (SIH) Key Discipline for Psychology in South China Normal University, and National Outstanding Youth Science Fund Project of National Natural Science Foundation of China (32022032).

## Author contributions

1. J. L., Conceptualization, Formal analysis, Investigation, Visualization, Methodology, Writing original draft; Y. C., Investigation, Visualization, Methodology, Writing original draft; X. Z., Conceptualization, Resources, Supervision, Funding acquisition, Investigation, Methodology, Writing original draft, Project administration.

## Ethics

All experiments were carried out in accordance with the Declaration of Helsinki. All participants provided written informed consent prior to the start of the experiment, which was approved by the Research Ethics Committee at South China Normal University (2021-04-01).

## Data and Code Availability

All analyses were conducted using custom code in MATLAB and the EEGlab toolbox for EEG data analysis. The code and processed data used for the final analyses are available at https://osf.io/34cex/. Raw data for this study can be requested from the Lead Contact, and the authors confirm that all reasonable requests will be fulfilled.

## Notes

### Competing Interest Statement

The authors have declared no competing interest.

### Summary of Updates

Corrected methodological flaw for cross-frequency coupling: We replaced the weighted phase lag index (wPLI) with the appropriate 1:2 cross-frequency phase synchrony (CFS) using the phase-locking value (PLV) following Siebenhuehner et al., 2020 (PLoS Biol) and Palva et al., 2005 (J Neurosci). Accordingly, all related text, figure legends, and references have been updated to reflect this change. Strengthened brain-behavior correlation analysis: To address the potential influence of outliers, we performed leave-one-out cross-validation (all iterations yielded positive r values, range: 0.183-0.497) and Spearman rank correlation (r = 0.13, p = 0.57). The original scatter plot was removed, and we now interpret this correlation as preliminary evidence that requires replication in larger samples.

